# ‘Who Goes There?’ Quantifying Scavenger-Carcass Activity at Mass Mortality Events

**DOI:** 10.64898/2026.07.03.735954

**Authors:** Katie A. Barton, Patrick B. Finnerty, Stefanie J. Bonat, Beatriz Martínez-López, Niraj Y. Meisuria, Thomas M. Newsome, Alison J. Peel, Justine A. Smith, Victoria J. Brookes

## Abstract

Mass mortality events (MMEs) create sudden pulses of carrion that can alter how vertebrate scavengers use carcass resources, including the frequency, duration, and timing of species–carcass contacts. These changes could have implications for pathogen transmission at the scavenger–carcass interface. We aimed to develop and apply a reproducible analytical framework for using camera-trap data to quantify transmission-relevant vertebrate activity at carcass sites under differing carrion biomass scenarios. We studied experimental carcass plots (single carcass ∼43 kg; ‘mass mortality’ plots [10 carcasses, >350 kg total]; 6 of each) in Australia’s alpine ecosystem. The framework integrated descriptive summaries (bipartite network analysis, Kaplan–Meier curves) and marked temporal point-process models to characterise structural and temporal dimensions of species–carcass activity. Mass mortality plots had greater overall visitation duration, occurring as sustained activity (50% of visitation event volume by day 17), compared with intense then rapidly declining activity at single carcasses (50% by day 8). Mass mortality plots also had higher predicted daily arrival probability and contact hours across most species, indicating an extended window for pathogen transmission. This framework provides empirically derived contact parameters for MME-related disease spread models using camera-trap data to identify potential transmission pathways at the scavenger–carcass interface.

## 2. INTRODUCTION

Climate change and other anthropogenic disturbances are reshaping ecological systems such that the annual likelihood of emerging infectious disease epidemics is predicted to increase in the coming decades (1, 2). Most EIDs arise from animal reservoirs and are associated with ecological “interfaces” where humans, domestic animals, wildlife and their associated microbial communities interact (3). While considerable attention has focused on interfaces such as live animal markets, agricultural systems, and wildlife-livestock boundaries, animal carcasses represent a comparatively underexplored interface (4). Carcasses are spatially discrete and temporally dynamic resources that concentrate scavenging activity and environmental microbial reservoirs, creating conditions conducive to microbial exchange among species (5–7). The ecological significance of the carcass interface is increasingly important because carcass availability in many ecosystems is rising, driven by both more frequent mass mortality events (MMEs) and declines in apex scavenger populations (8, 9). Therefore, predicting scavenger assemblages and scavenger-carcass contact dynamics in response to increases in carrion availability is important to understand the influence of MMEs and declines in apex scavenger populations on microbial transmission at the carcass interface.

MMEs are extreme influxes of carrion into ecosystems due to the sudden and substantial death of a large percentage of an animal population (10). Examples include the 2020 mass mortalities of African elephants (*Loxodonta africana*) in Botswana and Zimbabwe, attributed to *Pasteurella multocida*-induced haemorrhagic septicaemia, and Australian flying-foxes (*Pteropus poliocephalus*) during extreme heatwave events (11, 12). Such events can substantially increase the density and persistence of carcasses in the environment, extending the period over which pathogen exposure and transmission opportunities may occur (4). Microbial transmission from carrion can be indirect (for example, African swine fever virus (ASFV) can be detected in the soil around ASFV-infected wild boar (*Sus scrofa*) carcasses for approximately one month (13)), or direct via consumption of infectious necro-material (for example, tuberculosis, botulism, and highly pathogenic avian influenza (HPAI) (14–17)). Consequently, increases in carcass abundance and persistence have the potential to alter the carcass interface and create conditions that facilitate microbial transmission, pathogen maintenance in animal reservoirs, and ultimately, EIDs.

Concurrently, many global apex scavenger populations are declining due to habitat loss, food shortages, poisoning, and other anthropogenic influences (9, 18, 19). Apex scavengers efficiently locate and consume carrion, reducing microbial load in ecosystems (20, 21). Consequently, they limit the exposure of opportunistic scavengers to potentially infectious necro-material, and thus the opportunity for ongoing transmission through inter- and intra-specific congregation at carcasses (22–26). For example, declining vulture populations in India have contributed to an increase in scavenging dog populations with free-roaming dog congregation at carcasses, and thus, increased rabies incidence (27). Scavenger species can differ markedly in frequency, duration, and timing of interaction with carcasses, creating heterogeneous pathways for potential pathogen spread (28). Despite this, there remains no widely adopted, reproducible framework to quantify how increased carrion availability influences scavenger contact dynamics at carcasses in ways relevant to pathogen transmission processes.

We address this gap by developing an analytical framework that integrates camera-trap observations with scavenger–carcass bipartite networks and temporal contact metrics to describe species assemblages and quantify vertebrate contact dynamics at carcass sites. We apply this framework to experimentally elevated carrion availability in an Australian alpine system where there are no obligate vertebrate scavengers, relatively low or variable densities of apex scavengers (for example, dingoes [*Canis dingo*] and wedge-tailed eagles [*Aquila audax*]) and smaller mesoscavengers dominate carrion use (26). This system provides an informative analogue for scavenger-carcass contact dynamics under anthropogenically-driven ecological change (9). We hypothesised that elevated carrion availability would (i) attract a greater richness of species, (ii) support a greater number and duration of species visitation and scavenging events, and (iii) increase the probability of daily arrivals and cumulative contact time for all species. By quantifying how carrion pulses restructure vertebrate contact dynamics, our framework aims to provide parameters for ecological components of infectious disease models and improve understanding of how MMEs may influence pathogen transmission opportunities in changing ecosystems.

## 3. METHODS

### 3.1. DATA COLLECTION

An experimental study was conducted across three sites in Kosciuszko National Park, an alpine region in southeastern Australia (*Figure 1*). Sites 1 and 3 were in montane vegetation, and site 2 in a tableland basalt forest region. Site macroecological characteristics are described in *Table S1*. Each site contained four plots: two single carcass plots and two mass mortality plots (n = 10 carcasses), at least 75m apart (*Figure 1*). Carcasses were sourced from nearby management culls and allocated to both plots < 24 hours post-mortem. All single carcass plot were fallow deer (*Dama dama;* ∼43kg carrion). Mass mortality plots had both fallow deer and eastern grey kangaroo (*Macropus giganteus*) carcasses (equal mix of carcasses at each plot; 350-500kg carrion/plot).

**Figure 1:**
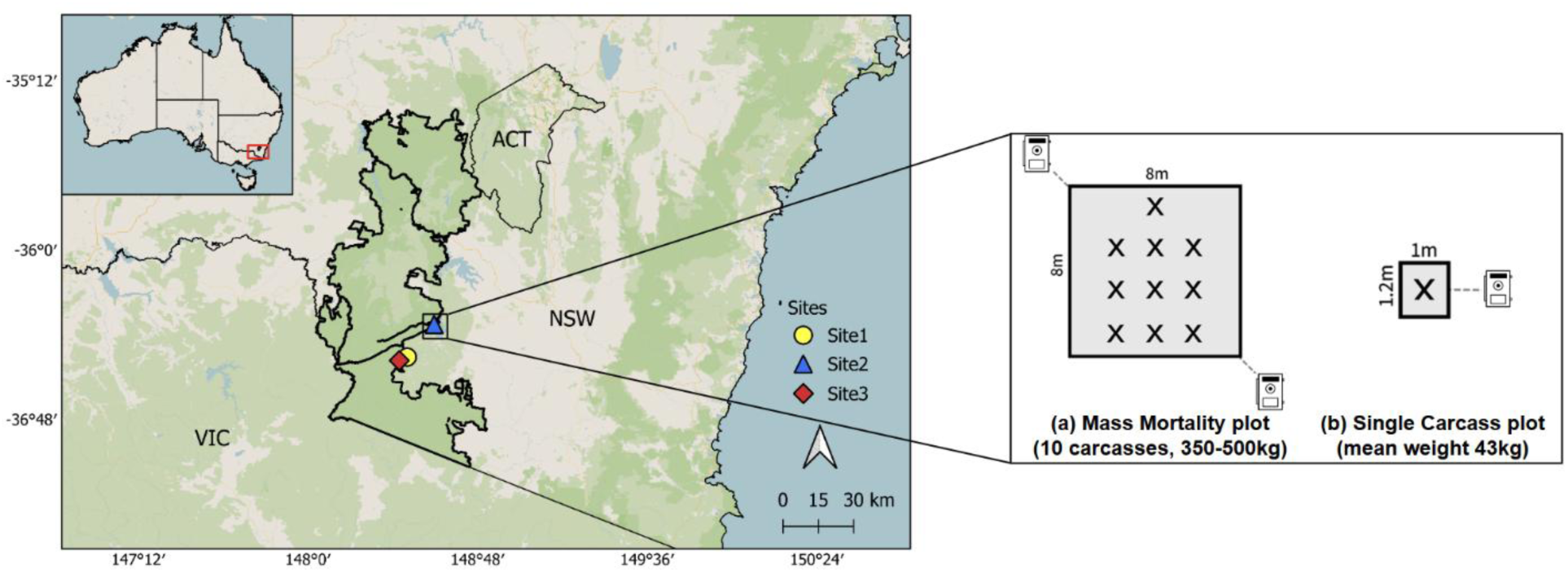
Study sites in Kosciuszko National Park, New South Wales (NSW), Australia. The inset map in the top left corner shows the relative location of the sites in Australia. The magnified inset of site 2 shows the carcass (crosses) and camera trap arrangement at each plot type. Each site had two of each plot arrangement (i.e. two of (a) and two of (b)). The map was generated with QGIS software version 3.4.0 (see Table S1 for spatial data source) (29).

Reconyx Hyperfire2 ™ camera traps (Reconyx Inc., Holmen, WI, USA) were used to identify vertebrate presence. At each single plot, one camera trap was positioned 1-2m of the carcass, approximately 1m above ground level. Two camera traps were positioned similarly at each mass mortality plot, in opposing corners to maximise vertebrate activity captured at the plot. Each trap was programmed to rapid-fire mode, which takes 10 consecutive photographs (∼1 image/s) with no recovery period when triggered by heat and motion. Each site was monitored for 30 days, with sites 1 and 2 monitored in December 2020, and site 3 in November 2021 (Australian summer).

### 3.2. DATA PROCESSING

Camera trap images were manually tagged with species present, group size, and behaviour (visiting or scavenging) using Digikam software (Version 7.6.0). Tagging was performed to the species-level, except for images of ravens and crows, which were all tagged as Corvidae (family-level) because they were challenging to distinguish morphologically due to their similar size and plumage colour (30). Images were grouped into visitation or scavenging ‘events.’ The classification of ‘visitation’ accounted for activity around carcasses that could be important for indirect microbial transmission. In events during which an animal fed (oral contact with a carcass) on at least one occasion, the whole event was tagged as scavenging for that species. Image metadata was extracted with the camtrapR package in R (31).

To determine the appropriate threshold of ‘event’ independence, a histogram was used to visualise the frequency of time differences between images of the same species, using 1-minute increments (32) (*Figure S1*). When time differences of <1 minute and >30 minutes were excluded, 85% of tagged images had <9 minutes between images; therefore, a new event was defined when there was ≥8 minutes since the last image of that species. This approach accounted for incidents when an animal was out of the camera trap field of view for a short time but in the vicinity of the carcass site. Images were pooled from both cameras at mass mortality plots; the use of ‘events’ in the analysis and not individual images minimised double counting of individuals (33).

### 3.3. DESCRIPTIVE ANALYSES OF SCAVENGER ACTIVITY STRUCTURE

All analyses were performed in R version 4.4.2 (34).

#### 3.3.1. Surveillance effort and species detections

Summary statistics were produced to describe surveillance effort, measured as camera-trap nights (24-hour recording periods), and broad species detection metrics, including abundance and group size.

#### 3.3.2. Descriptive network analysis

To describe total species activity associated with plot types, static egocentric bipartite networks were constructed from interaction matrices between vertebrate species (rows) and carcass plots (columns), with edges representing visitation events, weighted by total visit duration. Each network was visualised using Sankey plots (sankeyNetwork function from the ‘networkD3’ package; (35)), and all metrics were determined with the ‘bipartite’ package (36). The bipartite package, as opposed to other egocentric network packages, was used because it better reflects the lack of alter-alter ties in the dataset (i.e. only carcass-species interactions and rare occurrence of species-species interactions).

Degree centrality identified species richness at carcass plot type (single or mass mortality plot). Weighted nestedness based on overlap and decreasing fill (WNODF) determined nestedness (37). Modularity was calculated using Beckett’s weighted extension of Barber’s bipartite modularity equation to examine whether species-carcass interactions occurred in distinct ‘modules’ (38, 39) (*Table 1*).

**Table 1:** Definition of each network metric used in this study.

| **Metric** | **Definition and interpretation** |
| --- | --- |
| *Degree centrality* | The number of connections of a node (40). In the present study, this indicates the number of carcass plots visited (species node) or number of species which visited during the study period (carcass plot node). |
| *Weighted nestedness based on overlap and decreasing fill (WNODF)* | A quantitative extension of the binary NODF metric designed to measure nestedness, a structural pattern where nodes with smaller degrees (specialists) interact with a subset of the interaction partners of nodes with higher degrees (generalists) (37, 41). In egocentric bipartite networks, WNODF determines if the network links (e.g., species and the carcass site they attend) are structured such that less-connected species only engage in a subset of the activities of more-connected species. Values range from 0 (no nestedness) to 100 (perfect nestedness). |
| *Modularity* | Modularity identifies clusters of nodes with high densities of links amongst themselves. In this study, a high modularity would indicate that certain species tends towards activity at one carcass rather than all four carcass plots. Values range from 0 (no modules) to 1 (distinct modules) (42). |

To account for varying network connectance and enable comparison of network metrics between sites, statistical null models were implemented in which 1000 randomised variations of each empirical network were simulated. Following (43) and (44), the ‘vaznull’ method was used in which network connectance and matrix dimensions are fixed and the number of interactions by each species fluctuates with each randomised network (45). Network metrics were standardised across the sites (Equation 1) (46–48). A histogram of null model values was assessed for skewness (skewness function from the ‘e1071’ package; (49)) to reduce the Type I error. If the skewness value was ≥0.5 (absolute value), data were log-transformed (50).

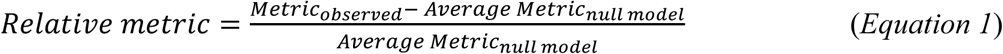

#### 3.3.3 Rate of visitation and scavenging activity

Kaplan-Meier (KM) curves were used to describe the rate of species activity (visitation and scavenging events) over time at each site. Curves were generated using the ‘survival’ and ‘survminer’ packages (51, 52). Events were weighted by duration (seconds) to account for plots with many events of short duration.

### 3.4. STATISTICAL MODELLING OF SPECIES-CARCASS ACTIVITY

To estimate daily species-specific species-carcass contacts, a marked temporal point-process model was implemented using a two-stage process: a generalised additive model (GAM) for scavenger arrival intensity and a GAM for visitation or scavenging duration (‘mgcv’ package, (53)). Sites were modelled individually, consistent with descriptive analyses and the diversity in macroecological variables between sites (*Table S1*).

For the intensity model, the response variable was event count/30-minute bin, fitted with a Poisson error distribution and a log-link. To ensure estimate event frequency rates (rather than counts), an offset of the natural logarithm of the bin duration (1800s) was used. For the duration model, the response variable was the logarithm of event duration(s)/30-minute bin, using a Gaussian error distribution and a log-link, with an offset structure applied as with the intensity model. For both models, temporal variation was modelled using cubic regression splines incorporating interaction terms between species and plot type. Based on recommendations for models with >100 data points, five knots were selected to balance over-fitting and the non-linear trend (54). Species and plot type were included as fixed parametric effects. All parameters were estimated using restricted maximum-likelihood (REML). Statistical significance was assessed at α = 0.05, and model fit was evaluated using overdispersion checks and standard residual diagnostics.

Using model outputs, an hourly prediction grid was constructed for each species, plot type, and event type for the 30-day period. Predictions for hourly arrival rate (*λ*_ℎ_) and mean event duration conditional on an event occurring (*μ*_ℎ_) were generated using the ‘predict’ function. As duration was modelled on the log scale, mean event duration on the original scale was estimated as exp(*μ* + *σ*^2^/2), in which *μ* is the predicted mean on the log scale and *σ^2^* is the residual variance (55). From these predictions, we calculated the instantaneous probability that a plot was occupied by at least one individual of a given species, the probability of at least one arrival within a 24-hour period, and the expected daily contact hours using Equations 2–4, and predictions were then visualised using ggplot2 (56).

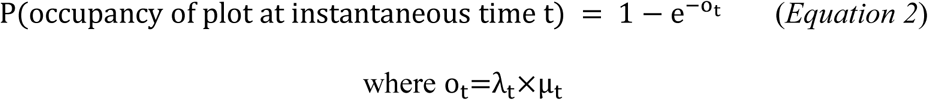

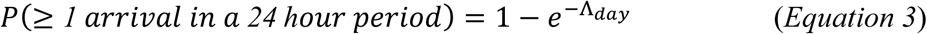

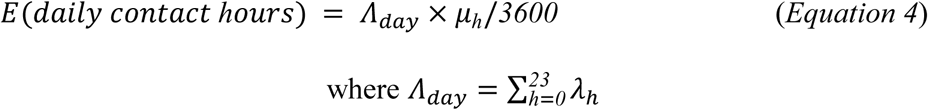

## 4. RESULTS

### 4.1. Surveillance effort and species detections

During the 30-day study period, 441,481 camera trap images contained identifiable species, comprising 2806 visitation and 1886 scavenging events across the three sites. Of a possible 720 trap nights (24 traps x 30 days), 688 trap nights were recorded (96%), with missing data due to camera malfunction or theft. Twenty-five vertebrate species were detected as visitors to carcass plots including eight scavengers: kookaburra (*Dacelo novaeguineae*), red fox (*Vulpes vulpes*), wedge-tailed eagle, fallow deer (*Dama dama*), common brushtail possum (*Trichosurus vulpecula*), domestic dog (*Canis familiaris*), dingo, and corvids (*Corvidae*) (*Figure 3*). The remaining species were: Australian magpie (*Gymnorhina tibicen*), blue-tongued lizard (*Tiliqua*), common wombat (*Vombatus ursinus*), crimson rosella (*Platycercus elegans*), mountain brushtail possum (*Trichosurus cunninghami*), red deer (*Cervus elaphus*), swamp wallaby (*Wallabia bicolor),* sambar deer (*Rusa unicolor),* white-winged chough (*Corcorax melanorhamphos),* eastern grey kangaroo (*Macropus giganteus)*, willie wagtail (*Rhipidura leucophrys)*, red-necked wallaby (*Notamacropus rufogriseus)*, rabbit (*Oryctolagus cuniculus*), horse (*Equus caballus)*, copperhead (*Austrelaps*), an unidentifiable bird, and an unidentifiable macropod (*Figure 2*).

**Figure 2:**
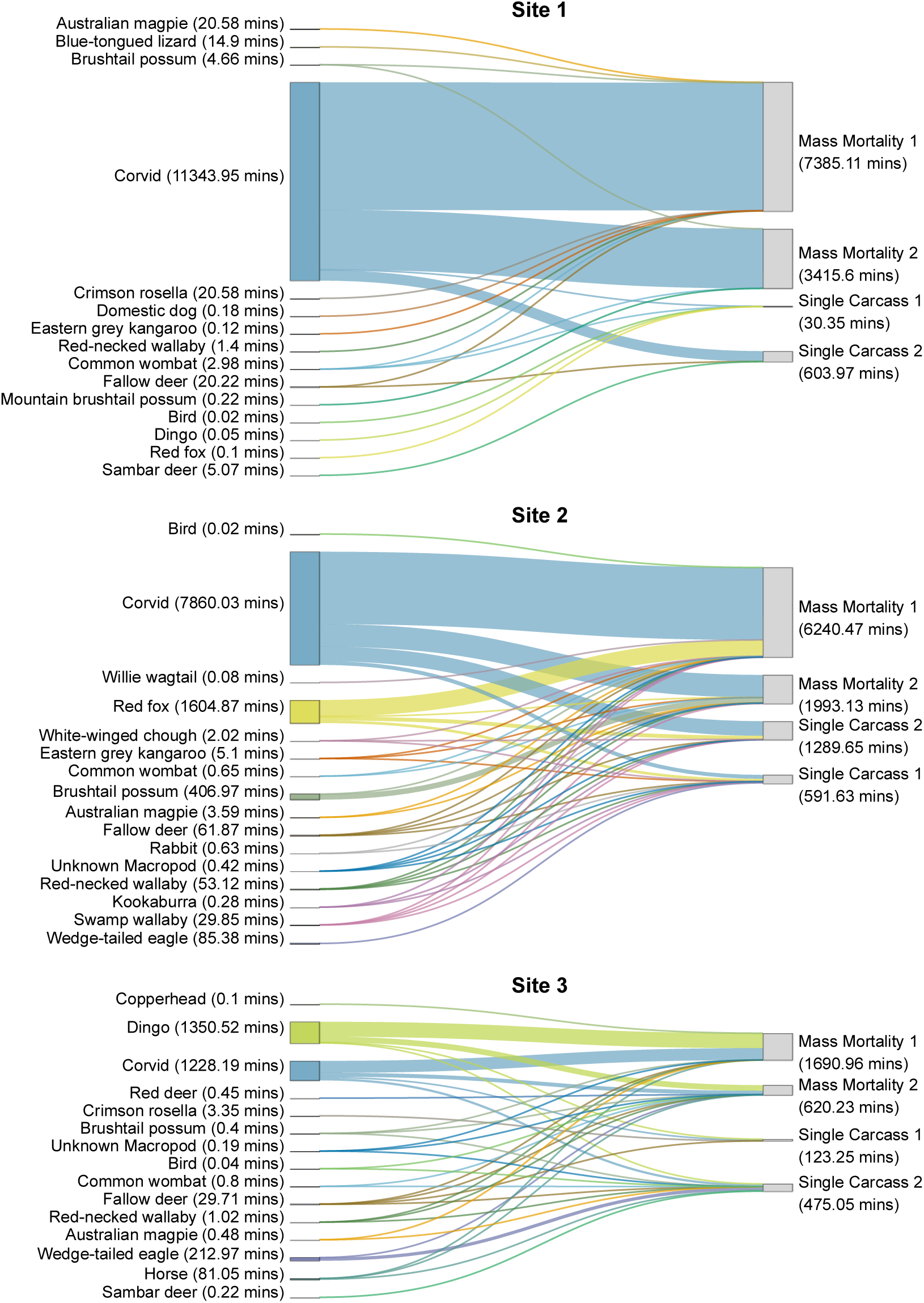
Egocentric bipartite networks of species and carcass plots during the 30-day study period at the three study sites. Nodes on the left represent species, nodes on the right represent carcass plots, and edges are weighted by the total visitation duration.

**Figure 3:**
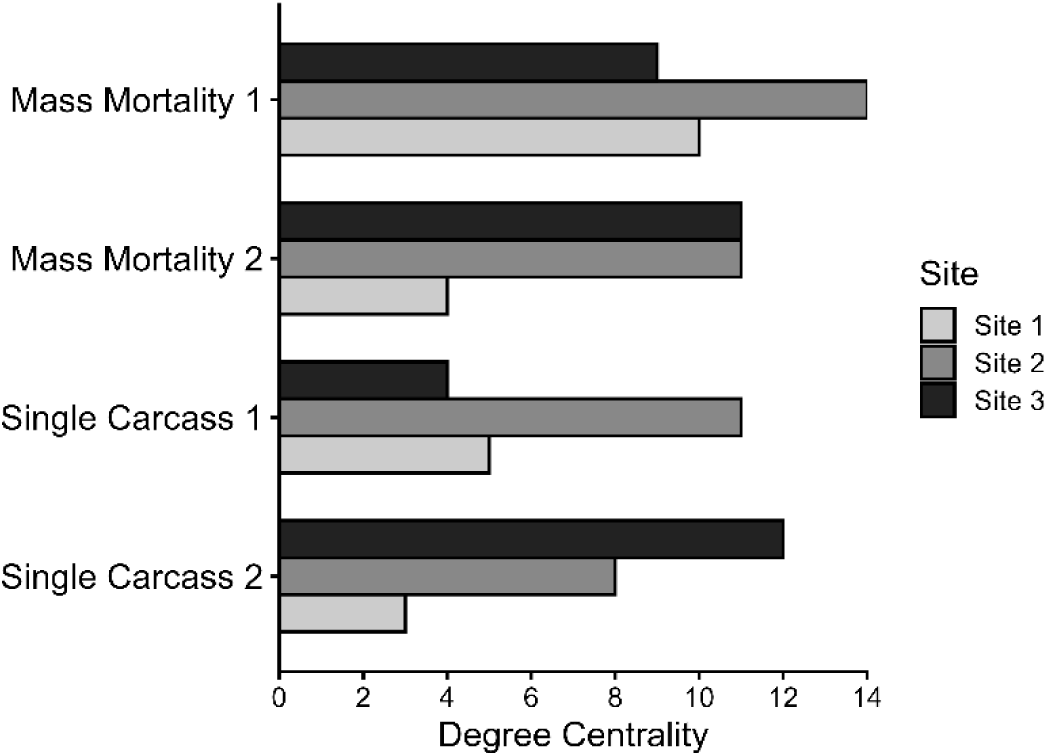
Bar plots of egocentric bipartite network degree by carcass plot and site.

The most frequent visitors were corvids, with 1802 visitation events across all sites (64% of total events), followed by dingoes (271 events, 9.7% but 270 of these were at site 3 only), and red foxes (268 events, 9.6%). Over 75% of visitation events at each site were to mass mortality plots. The most frequent feeders were corvids with 1396 scavenging events across all sites (74%), followed by dingoes (223 events, 11.8%; with all of these at site 3 only), and red foxes (178 events, 9.4%). Over 80% of the scavenging events at each site were at mass mortality plots. Median group size was 1 for the most frequently scavenging species at both plot types, except at mass mortality plots for corvids (median 2) with ranges of 1-7 (single plot) and 1-17 (mass mortality plot), for red foxes with ranges of 1-3 (single plot) and 1-4 (mass mortality plot), and dingoes (range 1-2 at both plot types).

### 4.2 Descriptive network analysis

Throughout all sites, degree centrality was not consistently greater at mass mortality plots than single plots and varied greatly both within and between sites. For example, the greatest degree centrality at sites 1 and 2 was at mass mortality plot 1 (degree centrality 10 and 14, respectively) and at site 3 was single plot 2 (degree centrality 12) (*Figure 3*). The total visitation duration was consistently greater at mass mortality plots than single carcass plots (despite a lower duration at mass mortality plot 2 compared to plot 1) (*Figure 2*).

At site 1, corvids were the only species to visit all four carcass plots (degree centrality 4, *Figure S2*). At site 2, swamp wallabies, red foxes, red-necked wallabies, unknown macropods, and fallow deer all had a degree centrality of 4. At site 3, dingoes, fallow deer, and corvids had a degree centrality of 4. The highest visitation duration was attributed to corvids at site 1 and 2, and dingoes at site 3.

Relative WNODF (rWNODF) was negative at all sites, indicating no significant nestedness. The relative modularity (rM; log transformed due to skewness >0.5) was negative at all sites, indicating no significant modularity (*Table 2*).

**Table 2:** Quantitative metrics calculated for the egocentric bipartite network for each site. Edges were weighted by visitation event duration (in minutes).

| Site | Metric | Observed | Null Mean | Null SD | Skewness | Log-scale | Z-Score | Relative measure |
| --- | --- | --- | --- | --- | --- | --- | --- | --- |
| 1 | Weighted NODF | 27.8 | 43.3 | 5.18 | -0.37 | N | -3.01 | -0.37 |
|  | Modularity | 0.05 | 0.002 | 0.0005 | 0.84 | Y | 3.8 | -0.13 |
| 2 | Weighted NODF | 48.5 | 59.9 | 4.08 | -0.23 | N | -2.79 | -0.19 |
|  | Modularity | 0.08 | 0.07 | 0.002 | 0.7 | Y | 8.17 | -0.58 |
| 3 | Weighted NODF | 47.8 | 54.9 | 4.05 | 0.16 | N | -1.76 | -0.13 |
|  | Modularity | 0.17 | 0.017 | 0.0048 | 0.63 | Y | 8.17 | -0.57 |

### 4.3. Rate of visitation and scavenging activity

Visitation at single plots primarily occurred early in the carcass deployment period, with the median time to 50% of total visitation event volume (number of events weighted by duration) at days 4, 4, and 8 at sites 1, 2, and 3 respectively (*Figure 4, a*). In comparison, mass mortality plots were visited at a consistent rate over the 30-day period, with median total visitation event volume at 12, 17, and 15 days at sites 1, 2, and 3 respectively. This general trend was also observed for scavenging events (*Figure 4, b*).

**Figure 4:**
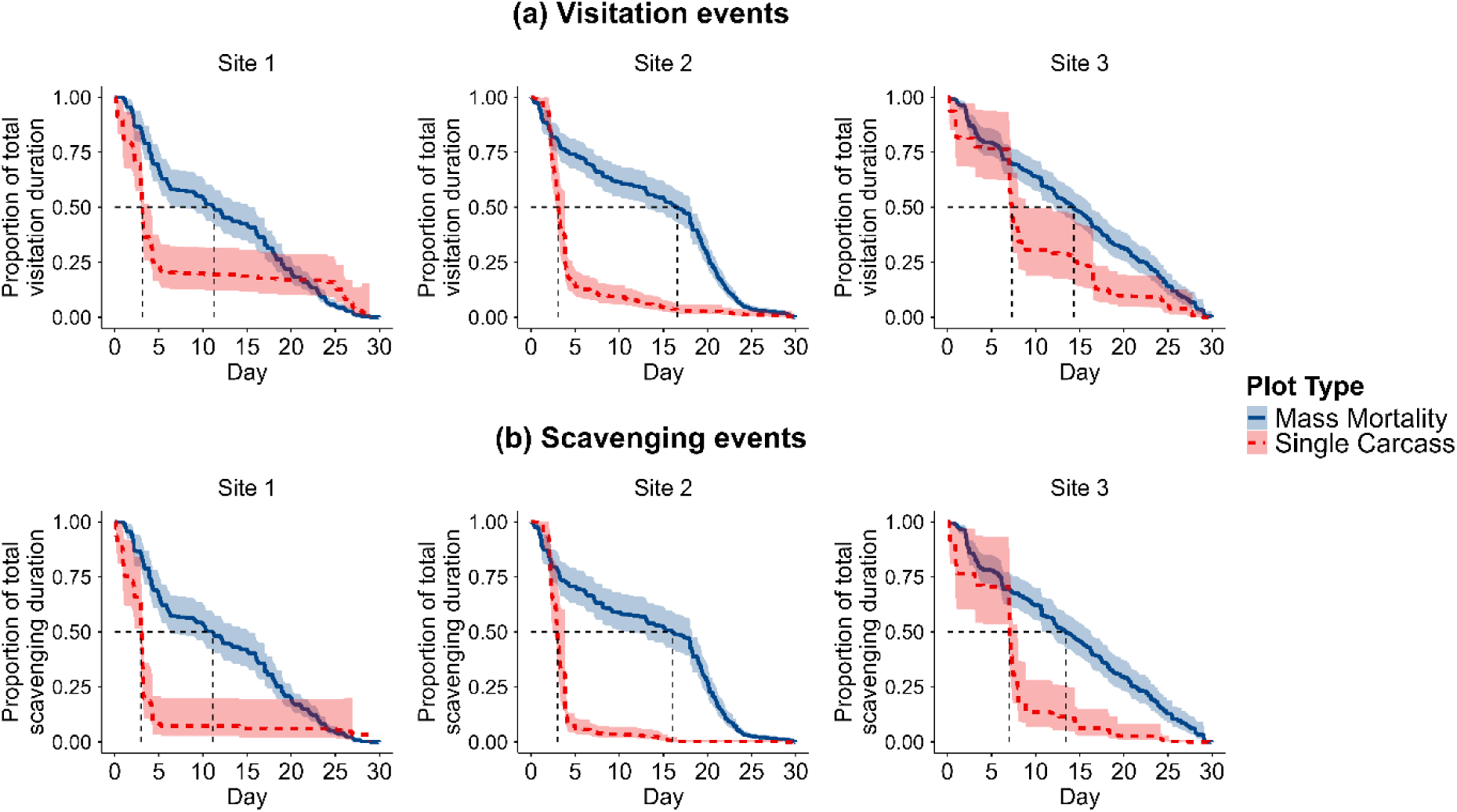
Kaplan-Meier curves for visitation events (a) and scavenging events (b), weighted by the duration in minutes. Shading indicates the 95% confidence interval. Dotted lines indicate median event occurrences.

### 4.4. Statistical modelling of species-carcass activity

Fourteen species had total visitation durations of ≥30 minutes during the 30-day study period and were therefore included in the visualisation of species-carcass activity. Although corvids had the highest predicted probability of ≥1 visitation/day at all sites throughout the study period, visitation probabilities differed between plot types (*Figure 5*). At mass mortality plots, the probability of ≥1 corvid visitation(s)/day was predicted to be 1.0 almost daily at all sites. In contrast, the predicted probability of ≥1 corvid visitation(s)/day at single carcass plots was initially high then declining throughout the study period at site 2, and declining to <0.4 at sites 1 and 3 mid-study before increasing again. A higher probability of ≥1 visitation(s)/day at mass mortality plots was consistent for all species except fallow deer and red foxes, which displayed an initially higher probability at site 2 single carcass plots.

**Figure 5:**
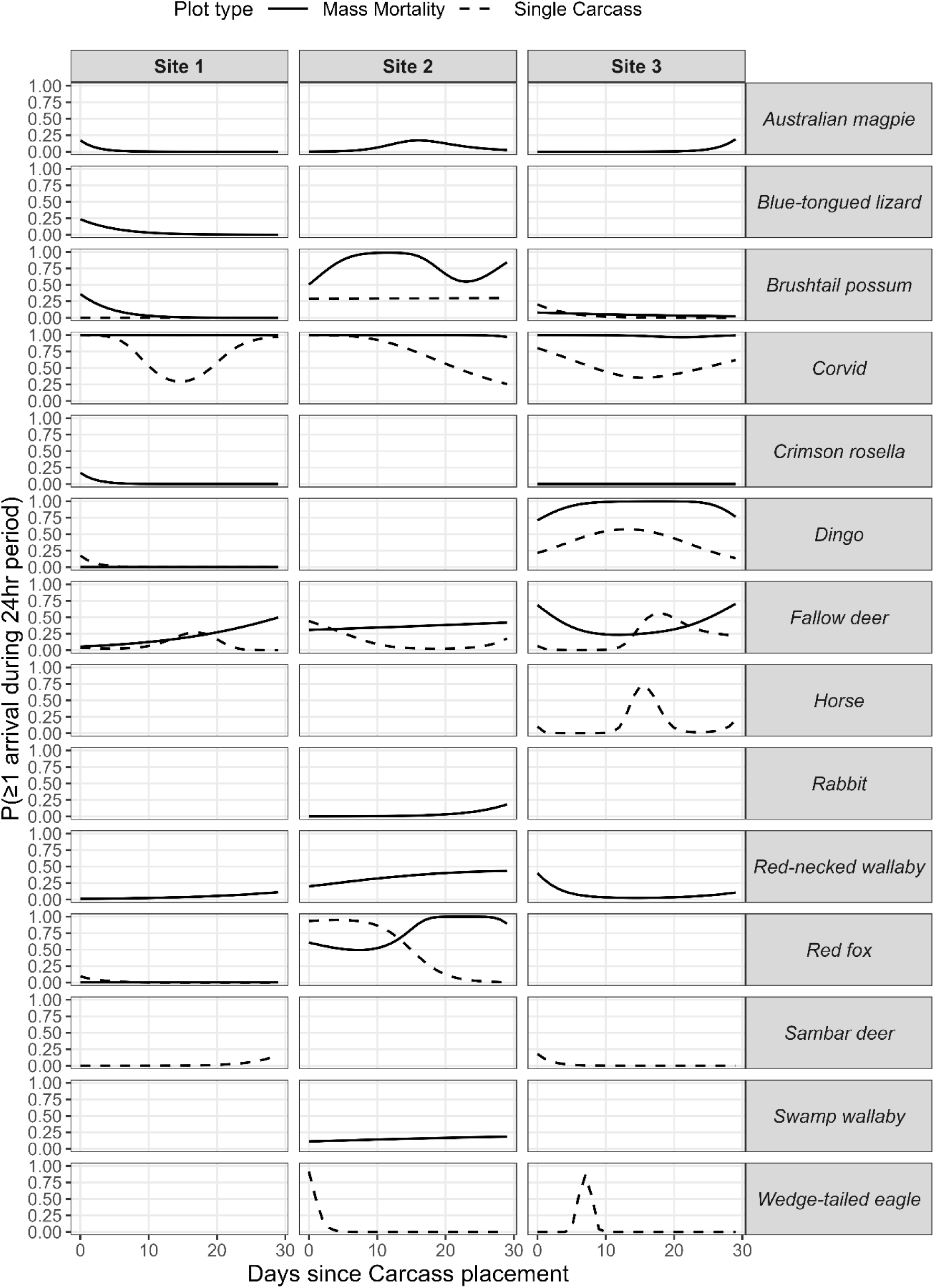
The predicted species-specific probability of ≥1 visitation event/24-hour period (day) across each carcass plot type at each site. Smooths were only plotted if the species was present for ≥30 minutes throughout the study period and were generated by the marked temporal point process (MTPP) model.

Five species had >30 minutes scavenging recorded throughout the study period. Similar to visitation, corvids were the most likely scavengers. The predicted probability of ≥1 scavenging corvid/day at mass mortality plots was consistently 1.0 for sites 1 and 2, with a decrease to approximately 0.8 at day 20, followed by a steady increase at site 3 (*Figure S3*). At single carcass plots, the predicted probability patterns of ≥1 scavenging corvid/day were highly variable: at site 1, the predicted probability decreased then increased, at site 2 it decreased monotonically, and at site 3 it decreased, increased, then decreased again (range 0.06 – 0.74). Dingoes only scavenged at site 3, and at both mass mortality and single carcass plots the predicted probability of ≥1 scavenging dingo/day increased then decreased with a predicted probability of 1 from 10-20 days. Consistent with visitation, a higher probability of ≥1 scavenging event/day at mass mortality plots was consistent for all species, although red foxes had an initially higher predicted probability of ≥1 scavenging event/day at site 2.

The predicted visitation hours/day were greater at mass mortality plots for each species throughout all sites, except for red foxes at site 2 (hours/day at single carcass plots were initially higher). Corvids had the maximum predicted visitation hours/day for each site, ranging up to 11.5 hours (*Figure 6*). Whilst predicted corvid visitation hours/day reduced over time, a bimodal trend was apparent at mass mortality plots at sites 1 and 2. For species that visited both mass mortality and single carcass plots, visitation hours/day were always greater at mass mortality plots, with the exceptions of red foxes at site 2, and fallow deer at site 3. Wedge-tailed eagles, rabbits, and horses each had 1-2 short periods of predicted visitation.

**Figure 6:**
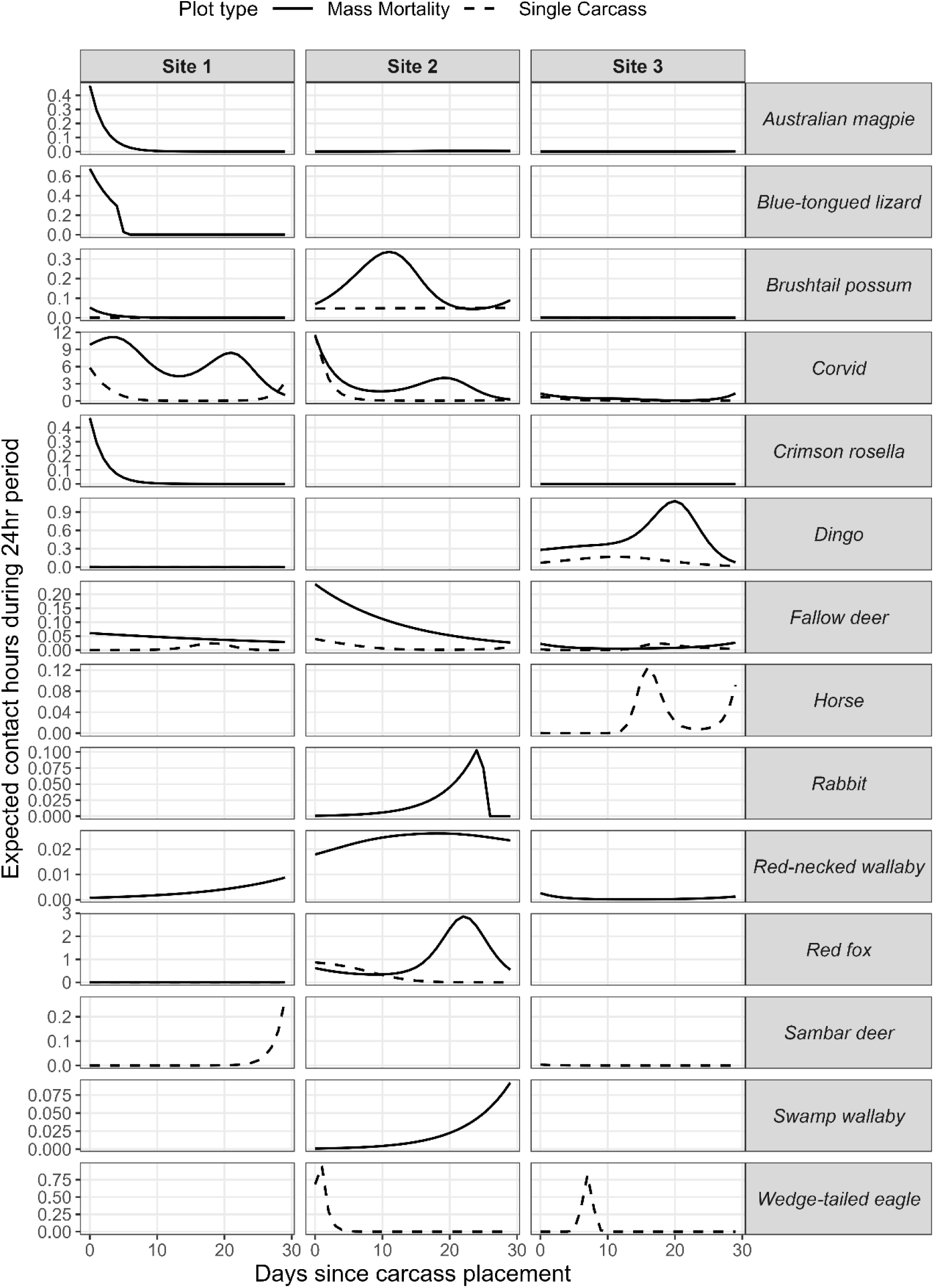
The predicted species-specific smooths for daily visitation contacts across the 30-day monitoring period across each carcass plot type at each site. Smooths were only plotted if the species was present for ≥30 minutes throughout the study period and were generated by the marked temporal point process (MTPP) model.

The predicted scavenging hours/day for corvids were initially higher at site 2 single carcass plots, before exponentially declining to a plateau before day 10 (*Figure S4*). This pattern was also evident for red foxes, with higher predicted contact hours at single carcass plots. However, by day 15, all species at all sites showed greater contact hours at mass mortality plots in comparison to single carcass plots. Notably, the scavenging hours of dingoes and red foxes at mass mortality plots initially declined between day 1 and 10 and was then followed by an increasing-decreasing trend (concave spline). Consistent with the visitation pattern, wedge-tailed eagles had a single peak in scavenging activity at site 2 and site 3, both of which occurred before day 10. The brushtail possum was only present scavenging for > 30 minutes at mass mortality plots, with varying patterns at each site.

Plots for predicted probability of occupancy/day are shown for visitation and scavenging events in *Figure S5* and *S6*. The shapes of these probability distributions are consistent with those in the total contact hours plots (visitation and scavenging).

## 5. DISCUSSION

Our reproducible framework for describing vertebrate activity dynamics relevant to microbial transmission at the carcass interface integrates complementary analytical steps: network summaries to characterise the structure and distribution of species–carcass interactions, Kaplan–Meier curves to describe the rate of visitation and feeding activity, and marked temporal point-process (MTPP) models to quantify species-specific contact probabilities and durations over time. Under experimentally increased carrion, assemblage structure and species–carcass contact patterns shifted toward greater and more sustained visitation and scavenging activity with higher predicted probabilities and durations of daily contact for most species, but without evidence of increased species richness or abundance. Site-level variation was substantial, indicating that comparative application across more systems will be needed to determine whether these patterns are generalisable and which ecological factors drive them.

Species richness, measured here as plot-level degree centrality, did not consistently differ between mass mortality and single carcass plots despite the substantially greater carrion biomass at mass mortality plots. This was not consistent with our first hypothesis and contrasts with studies from other systems in which larger carrion resources supported richer vertebrate scavenger guilds (57–59). Vertebrate species richness is important in the context of microbial spread because it is correlated with carcass removal rates, and can influence the extent of inter- and intra-specific activity around carcasses (60–62). Differences among sites, including habitat and surrounding anthropogenic features, could have contributed to this heterogeneity, consistent with work showing that scavenger guild structure can vary with human influence and landscape (63). Importantly, degree centrality is insensitive to whether activity was concentrated or sustained over time, and such temporal dimension is needed to determine microbial transmission opportunity (64, 65).

Total activity durations and quantitative metrics provided greater insight into differences in network structure between plot types. Across all sites, mass mortality plots consistently accumulated higher visitation duration and event frequency, indicating that visiting scavengers may be more likely exposed to pathogenic material at these sites compared to single carcass plots. This pattern aligns with expectations and previous studies showing greater scavenger activity at larger carrion resources in systems such as the South African savanna and Southwestern Montana (57, 66). Therefore, mass mortality nodes can be interpreted as the focal points of each site’s activity and thus represent greatest potential for microbial exposure.

The broader structural organisation of contacts was weak, limiting inference from network structure alone and emphasising the need for temporal analysis. Interpretation of key nodes depends on modularity and nestedness metrics (67). Modularity can restrict transmission to clustered groups of dense contacts, whereas nestedness can increase overlap in interactions and thus transmission potential across the scavenger guild (68). In this study, however, there was no significant modularity or nestedness, possibly due to the proximity of carcass plots and consistent with previous discussion of limitations of nestedness metrics in scavenger–carcass systems (63). As such, network metrics are most informative as a structural baseline identifying where contacts are concentrated, but do not fully capture how transmission risk evolves. Temporal trends in species-specific contacts are therefore critical for understanding how the risk profile of microbial transmission changes over time (69).

The marked temporal point-process (MTPP) models demonstrated a consistently higher predicted probability of daily species–carcass activity across nearly all species at mass mortality plots. Consistent with the overall rate of events at plots indicated by the Kaplan-Meier survival curves, this indicates that mass mortality plots exceeded the threshold of carrion that the local scavenger population. Corvids, a species identified as central in the network analyses, showed sustained probabilities of daily arrival at mass plots. This is consistent with findings from other Australian scavenger guild studies (32) and has important implications for disease transmission because scavenging corvids have been shown to disseminate pathogens via defaecation (70, 71) and to become infected through consumption of infected carcasses, including in the context of HPAI (72).

Mass mortality plots also showed more complex temporal patterns of contact, with evidence of two-phase activity compared to the single early pulse observed at single carcass plots. This suggests a shorter window of potential pathogen exposure at single carcasses and a more prolonged, potentially multi-phase window at mass mortality plots. While greater activity at mass mortality plots aligns with increased resource availability, the timing of peaks may reflect preferences for different stages of carcass decay. Similar patterns have been observed in corvids, where scavenging activity peaks during early and late decay phases, potentially linked to invertebrate dynamics (73). However, whether later peaks in contact translate into increased transmission risk will depend on pathogen persistence (22, 74). For environmentally persistent pathogens such as African swine fever virus or *Bacillus anthracis*, extended or secondary contact peaks may remain epidemiologically relevant (75–77).

Several limitations should be considered when interpreting these findings. Networks were constructed at the species level and with a limited number of carcass plots, which may restrict inference from network metrics (78). In addition, the GAM-based MTPP approach assumes independence between events and is intended to describe contact patterns rather than infer causal drivers of interactions. As such, the analysis captures species–carcass contact at a focal site, including temporally proximate use relevant to indirect transmission. Despite these constraints, the observed heterogeneity in species-specific scavenger–carcass contacts between single and mass mortality plots is informative for parameterising transmission processes, particularly the effective contact rate (β) in disease spread models, and helps reduce bias associated with relying solely on contact intensity or duration (79–81).

## 6. CONCLUSION

By integrating network summaries, time-to-activity curves, and species-specific temporal predictions, our framework quantifies carcass-interface contact opportunities, and captures complementary structural and dynamic dimensions of carcass use that would not be identified using a single analytical approach. These results provide empirically derived parameters for disease spread models, particularly the effective contact rate (β), and support the use of camera-trap data to characterise transmission-relevant contact processes in changing ecosystems. In the context of this study, increased carrion availability led to greater and more persistent species–carcass contacts, extending the window of potential pathogen transmission at mass mortality plots compared to single carcasses. Corvids dominated visitation and feeding activity across sites, suggesting a potentially important role in transmission pathways.

