## Supplementary material for "‘Who Goes There?’ Quantifying Scavenger-Carcass Activity at Mass Mortality Events"

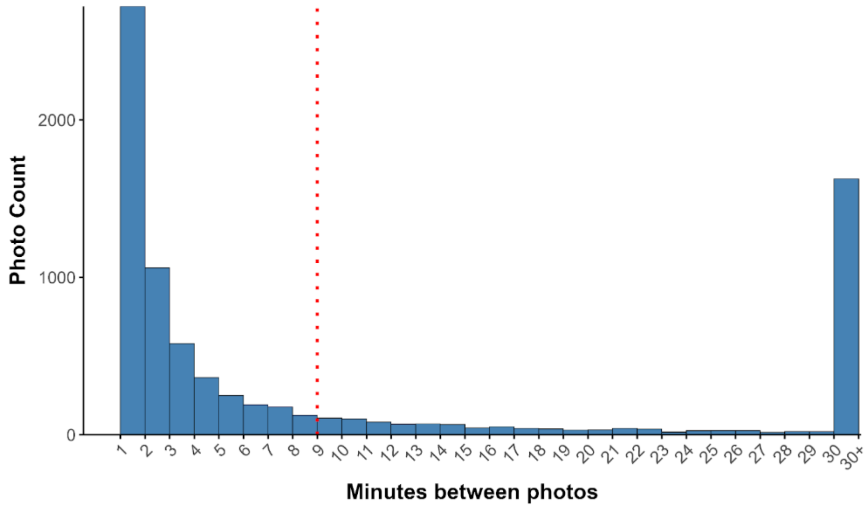


***Figure S1***: *Histogram of time differences between tagged photos of each species, based on* $\geq$*1-minute increments. The red dotted line indicates that 85% of photos were separated by less than 9 minutes, thus justifying the appropriateness of an 8-minute event threshold.*

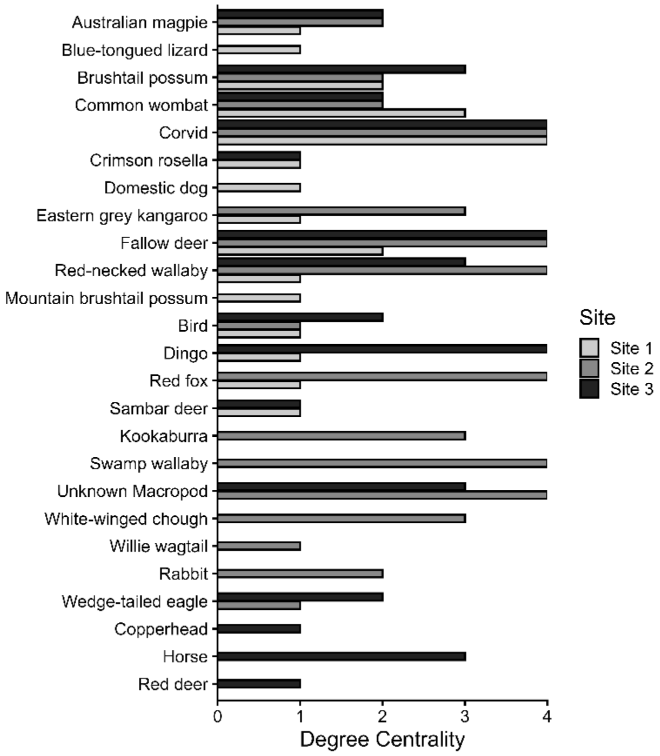


***Figure S2:*** *Bar plots of static bipartite network degree of all visiting vertebrate species and site. The first eight species to appear on the y-axis are those that scavenged throughout the study period.*

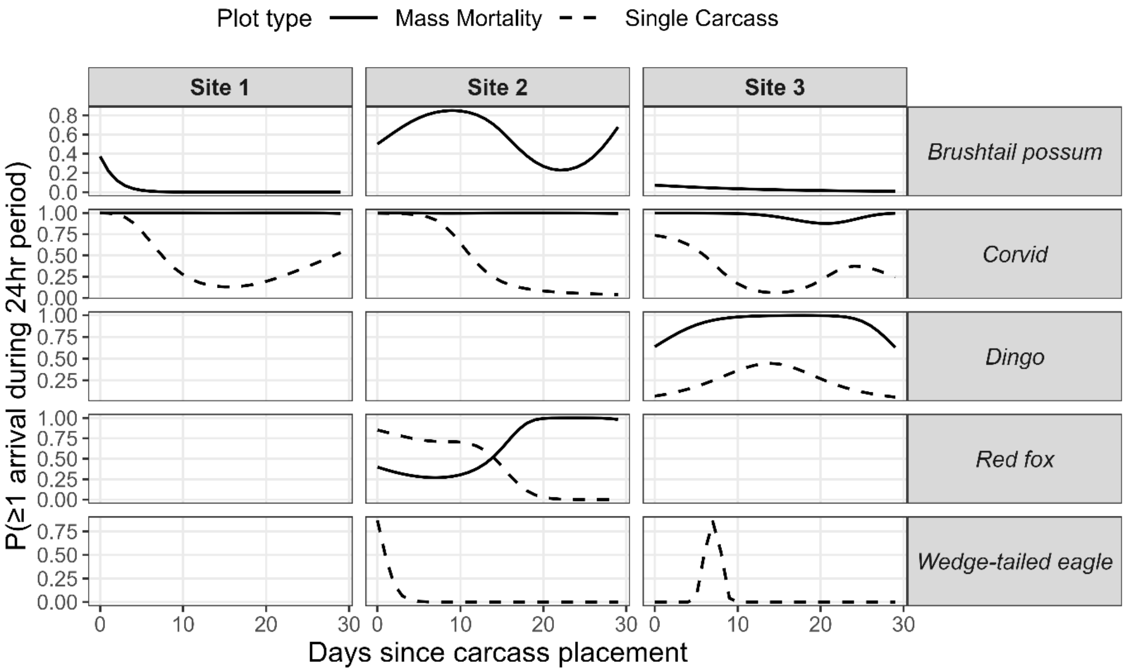


***Figure S3:*** *The predicted species-specific probability of* $\geq$*1 feeding event/24-hour period (day) across each carcass plot type at each site. Smooths were only plotted if the species was present for* $\geq$*30 minutes throughout the study period and were generated by the marked temporal point process (MTPP) model.*

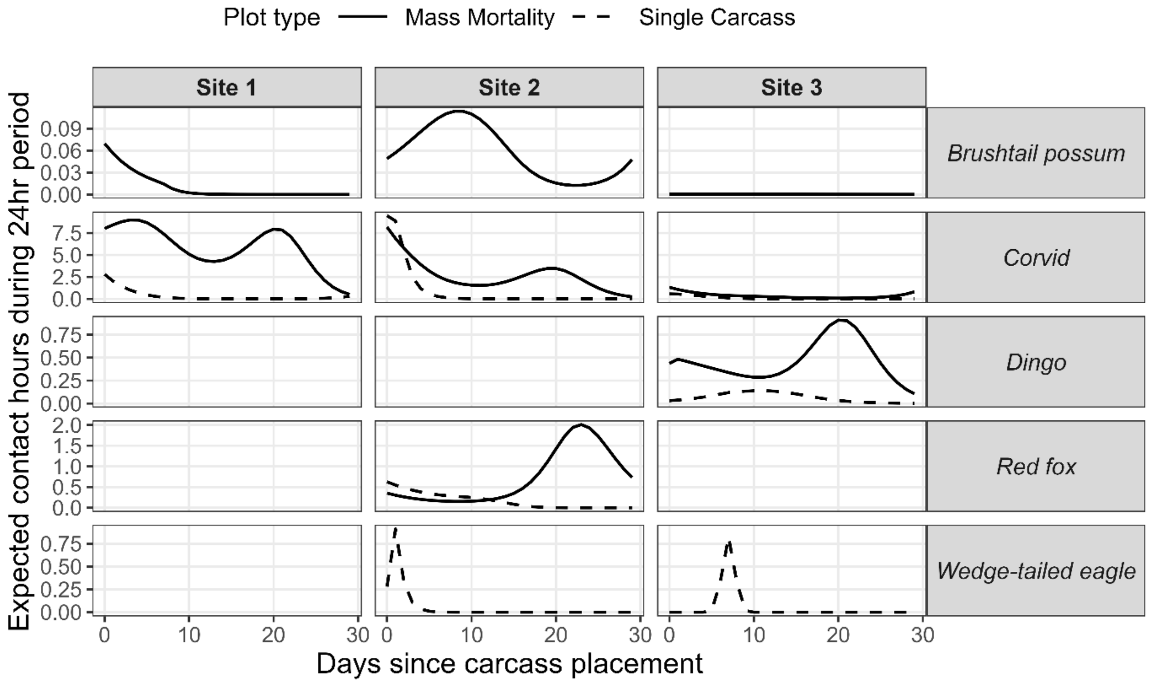


***Figure S4:*** *The predicted species-specific smooths for daily scavenger contacts across the 30-day monitoring period across each carcass plot type at each site. Smooths were only plotted if the species was present for* $\geq$*30 minutes throughout the study period and were generated by the marked temporal point process (MTPP) model.*

***Table S1:*** *Climatic and spatial variables for each carcass plot at each of the three sites. Using spatial coordinates, climatic and macroecological variables were extracted, the latter using the 'sf' package. All climatic variables were sourced from (data:* https://open-meteo.com/en/docs/historical-weather-api*, date accessed: 16th September 2025). The following spatial features were used: distance to the nearest water (data:* https://data.humdata.org/dataset/hotosm_aus_waterways*, date accessed: 7th September 2025), distance to the nearest road (data:* https://opendata.transport.nsw.gov.au/dataset/road-segment-data-from-datansw, date accessed: 6th September 2025*) and distance to the nearest farm (data:* https://datasets.seed.nsw.gov.au/dataset/nsw-landuse-2017-v1p5-f0ed-clone-a95d, date accessed: 7th September 2025*)).*

| Site | Carcass plot | Altitude (m a.s.l) | Distance to nearest road (m) | Distance to nearest farm (m) | Distance to nearest waterway (m) | Max daily temperature (°C) | Max daily precipitation total (mm) | Total precipitation (mm) | Average daily relative humidity (%) |
|---|---|---|---|---|---|---|---|---|---|
| 1 | *Mass Mortality 1* | 1131 | 426 | 375 | 77 | 24.6 | 28.8 | 106.5 | 75.2 |
| | *Mass Mortality 2* | 1137 | 505 | 533 | 177 | 27.6 | 28.8 | 110.1 | 73.5 |
| | *Single Carcass 1* | 1124 | 368 | 305 | 31 | 27.7 | 28.8 | 110.1 | 73.5 |
| | *Single Carcass 2* | 1164 | 625 | 513 | 281 | 27.4 | 28.8 | 110.1 | 73.5 |
| 2 | *Mass Mortality 1* | 976 | 291 | 1653 | 316 | 28.7 | 19.1 | 77.6 | 71.0 |
| | *Mass Mortality 2* | 970 | 198 | 1669 | 273 | 28.7 | 19.1 | 77.6 | 71.0 |
| | *Single Carcass 1* | 981 | 238 | 1853 | 604 | 31.2 | 23.6 | 95.6 | 70.6 |
| | *Single Carcass 2* | 976 | 183 | 1835 | 549 | 30.0 | 23.6 | 98.7 | 74.1 |
| 3 | *Mass Mortality 1* | 1172 | 2963 | 2996 | 105 | 25.0 | 82.2 | 233.9 | 78.3 |
| | *Mass Mortality 2* | 1167 | 2872 | 2986 | 65 | 25.0 | 82.2 | 233.9 | 78.3 |
| | *Single Carcass 1* | 1166 | 2771 | 2948 | 112 | 25.0 | 82.2 | 250.2 | 78.6 |
| | *Single Carcass 2* | 1158 | 2719 | 2967 | 84 | 25.1 | 82.2 | 238.2 | 77.2 |

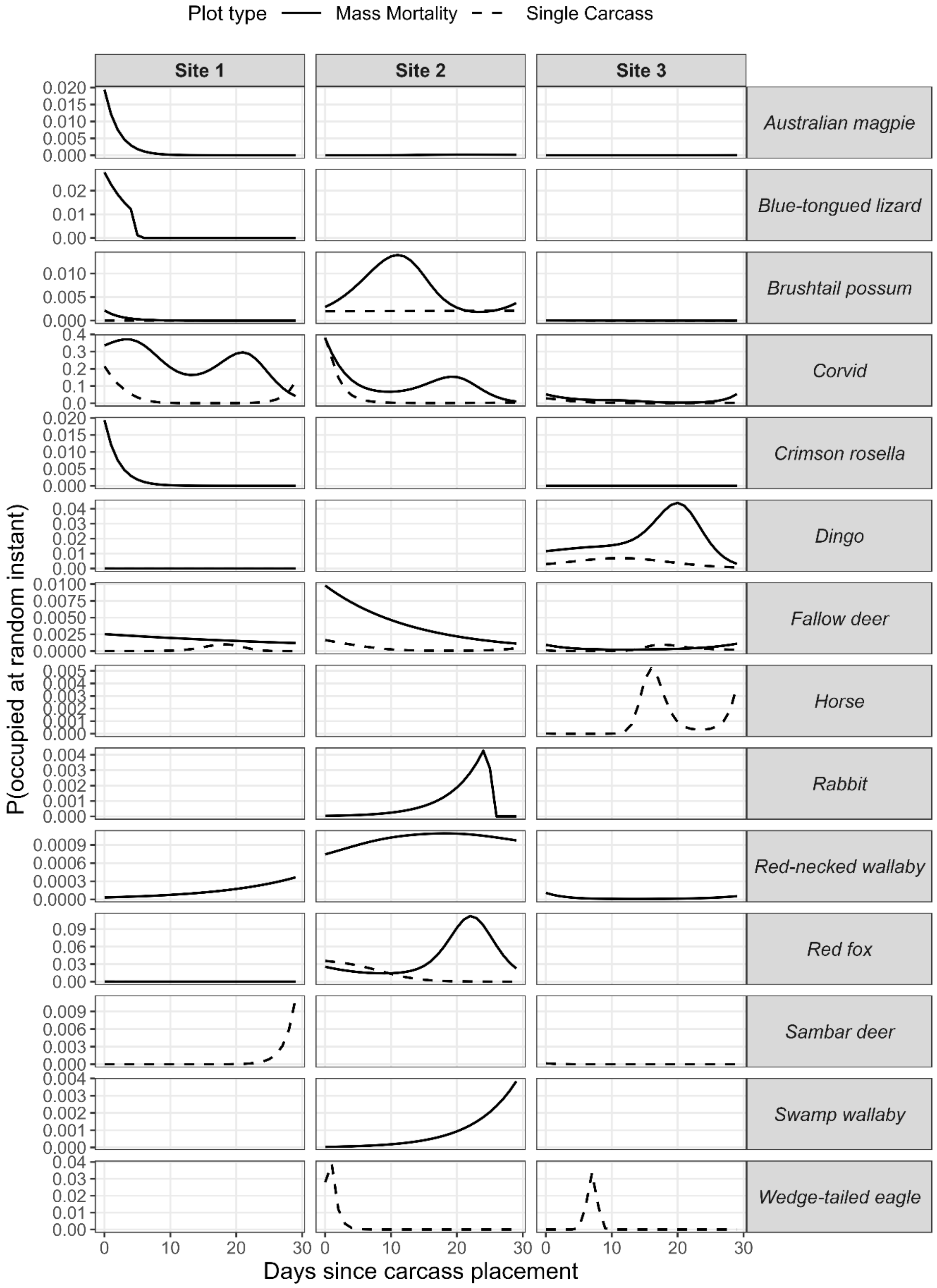


***Figure S5:*** *The predicted smooths for the instantaneous occupancy probability that a visiting species is present at each plot type (single carcass and mass mortality) at each site across the 30-day monitoring period. Smooths were only plotted if the species was present for ≥30 minutes throughout the study period and were generated by the marked temporal point process (MTPP) model.*

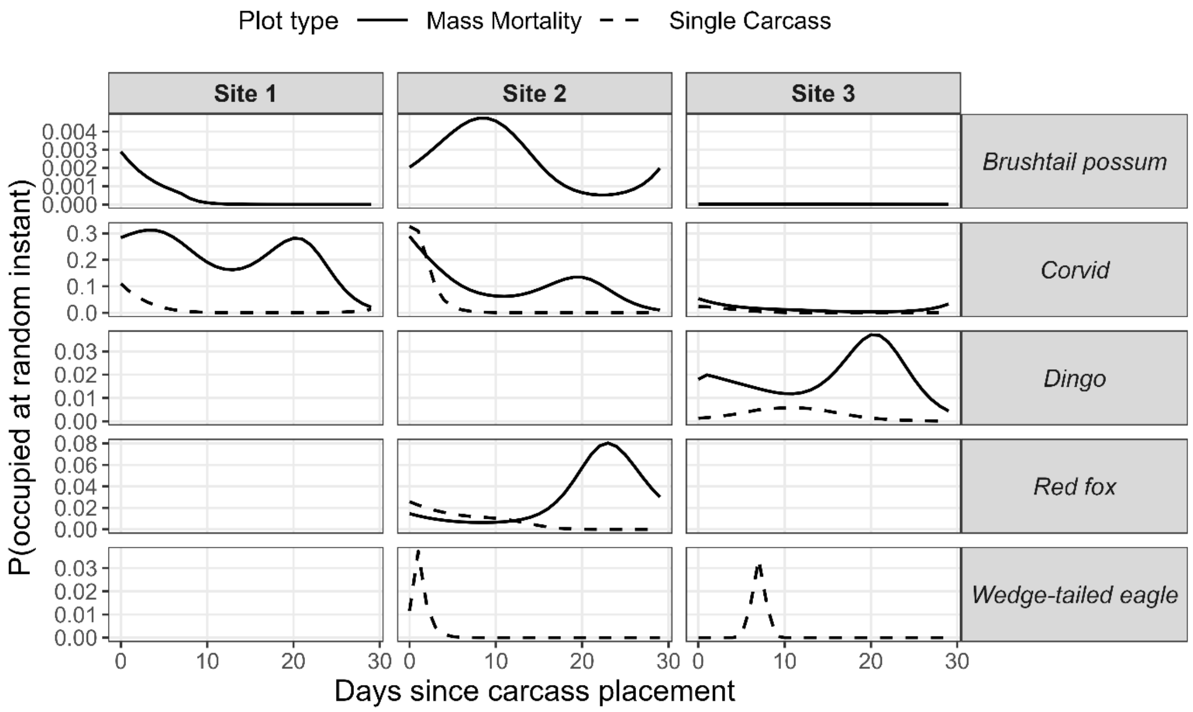


***Figure S6:*** *The predicted smooths for the instantaneous occupancy probability that a scavenging species is present at each plot type (single carcass and mass mortality) at each site across the 30-day monitoring period. Smooths were only plotted if the species was present for ≥30 minutes throughout the study period and were generated by the marked temporal point process (MTPP) model.*
